# Mucosal washes are useful for sampling intestinal mucus-associated microbiota despite low biomass

**DOI:** 10.1101/2023.12.12.571228

**Authors:** Jennifer N Martinez-Medina, Fereshteh Ghazisaeedi, Catharina Kramer, Jörn F Ziegler, Victoria McParland, Britta Siegmund, Víctor Hugo Jarquín-Díaz, Marcus Fulde, Sofia K Forslund

## Abstract

Exploring the dynamic relationship between mucus-associated microbiota and host health is pivotal, yet prevalent studies using stool samples may not accurately represent these bacteria. Here, we explored mucus-associated microbiota in the gastrointestinal tract of mice and the terminal ileum in humans, using three different sample types: mucosal washes, scraping, and intestinal content in mice and biopsies and mucosal washes in humans. We employed DNA quantification and 16S rRNA sequencing to assess how comparable the information yielded from different sample types, evaluating findings relative to expectations from state-of-the-art and under controlled benchmarks.

Mucosal washes in mice exhibited higher bacterial DNA and lower host DNA contamination than scraping samples. Similarly, in humans, washes surpassed biopsies in bacterial yield. Despite variations in read counts, microbiota diversity and composition remained remarkably similar between methods in both species, faithfully reflecting expected genotypic and phenotypic differences.

We conclude that washes reduce host contamination without inducing substantial compositional bias when sampling mucosal microbiota. Our findings emphasize mucosal washes as alternatives to biopsies in humans and scrapings in mice, providing insights for improving result transferability across hosts. Our research underscores the importance of considering the mucus-associated microbiota to track host-microbiome interactions closer to their actual interface surface.

## Background

Assessing taxonomic composition and functional profiles of intestinal microbiomes yield relevant insights into mechanisms and determinants of health and disease of their hosts.^1–3^ The well-being of that host depends in part on the balanced interaction with the microbiota throughout life and development, just as the microbiota depends on its host. In general, disruption of this balance promotes the development of chronic diseases, such as inflammatory bowel diseases.^4^ Most studies to date have focused on the fecal microbiota, which only imperfectly correlates as regards either role or composition with the microbial communities found in the intestinal mucus, or the directly mucus-associated microbiota which are a much tighter physiological relations to the host.^5^

Epithelial mucus is a glycoprotein secreted by the goblet cells that forms a layer covering the intestinal epithelium and provides a barrier between the lumen and the underlying tissue.^6^ The properties and composition of this mucus are essential for establishing and maintaining its associated microbiota.^7–9^ Proper characterization of the microbial communities inhabiting the mucus layer relies on the use of an appropriate sampling methodology that is representative of the biology under study.

In the mouse models commonly used to study intestinal diseases, the samples selected to study the mucus-associated microbiota are mainly obtained from luminal contents, or using scraping, washing, and from whole tissues upon sacrifice. In contrast, in humans, the most commonly employed sample types for the same purpose are biopsies, mucosal-luminal interface aspirates, colonic lavages, and endoscopic brushes.^10–13^ Mucosal biopsies are considered the gold standard sample for characterizing the human mucus-associated microbiota.^10^ However, they have several drawbacks, including: 1) a reduced representation of the sampled intestinal subsegment, and 2) high content of host DNA which may interfere with the bacterial DNA signal and further interpretation of its results, especially as the mucosal interface itself, in contrast with the luminal contents, is a sample with low bacterial biomass.^10,14^ Moreover, it is important to identify a protocol that works well under both clinical and preclinical conditions so that results from these modalities remain comparable, yet reflecting the constraints of these settings, including limitations from the sampling of patients as well the size from small model organisms like mice.

Body sites and samples with low bacterial biomass pose a major challenge for the study of mucus-associated microbial communities. Several attempts have been made to improve the assessment of mucus-associated microbiota. One approach is to deplete host DNA from biopsies by using different extraction methods to enrich for bacterial DNA prior to sequencing.^14^ Other alternatives involve different sample types, such as lavages, for reproducible substitution of mucosal biopsies.^10,11^ Furthermore, the variation in the sample type used within even the same host could pose technical and biological challenges for the interpretation of the results and the comparison between studies. Nevertheless, there is no consensus on the best choice of sampling methodology for the assessment of mucus-associated microbiota in humans or animal models.^5,10,15^

A critical step towards better characterization of the mucus-associated microbiota is to evaluate the reliability of low biomass samples such as biopsies, scrapings, or mucosal washes and compare them to luminal content from the small and large intestine of different host origins. Such benchmarking is necessary to assess the differences between samples obtained by different methods and between samples from different anatomical subsegments. This may help clarify the advantages and limitations of the respective techniques and provide a basis for possible large-scale study implementations. Here, we aim to compare the mucus-associated microbiota in different sample types collected from three gastrointestinal subsegments in mice, and from the terminal ileum in humans. We do this as a way to identify a protocol to allow for clinical-preclinical comparisons and improved reproducibility. This is essential in order to obtain a more robust and translatable understanding of the health implications of variation in the intestinal mucus-associated microbiome.

## Methodology

### Study sample

All animal experiments were approved by the local office of occupational health and technical safety “Landesamt für Gesundheit und Soziales, Berlin” LaGeSo Reg. Nr. T 0284/15. Animals were maintained in the Institute of Microbiology and Epizootics, School of Veterinary Medicine at the Freie Universität Berlin, Robert-von-Ostertag-Str. 7 14163 Berlin, Germany. We collected samples from C57BL/6J Toll-like receptor 5 null allele (B6(Cg)Tlr5<tm1.2Gewr>/J - 028909) (TLR5-KO) (N = 3) and wild-type (WT) (N = 3) mice, each of 8 weeks of age per genotype. These mice were under a maintenance diet for this study. The mutant genotype was selected as previous work showed it to exhibit phenotypes where alterations to the mucosal microbiome are expected.^16^

Human samples were obtained at the Department of Gastroenterology, Infectious Diseases and Rheumatology, Charité - Universitätsmedizin Berlin, Berlin, Germany between 2021 and 2022, and approval from the ethical committee was obtained. Biopsies were taken from 17 control and 19 Crohn’s disease (CD) patients, as part of an already scheduled colonoscopy and written informed consent was obtained. For the control group, indications for the exploratory colonoscopy were symptoms like abdominal pain, diarrhea, weight loss, surveillance, or endometriosis. For the patients with CD, colonoscopy was performed for assessing disease activity, surveillance, or specific symptoms. Patients in all stages of the disease were included (before or under treatment, and remission), while patients were excluded if they had undergone an ileocecal resection or an increased procedural risk in colonoscopy, like due to cardiopulmonary comorbidities or to the intake of anticoagulants. Although we included CD patients, it is not the purpose of this study to determine differences between healthy or disease groups. A higher sample size would be required to reach statistical power for epidemiological conclusions, and the present study was also intended as a methodological pilot to design such a study at a later stage.

### Collection of low-biomass sample types and intestinal content samples

We collected mucosal washes and scraping samples from the ileum, cecum, and colon of mice. The intestinal contents from the same subsegments were also collected as a reference for the corresponding luminal microbiome. We removed the intestines from the mice, separated the three subsegments of interest and opened them longitudinally. The ileum corresponded to the last third of the small intestine in mice, while the colon corresponded to the first third of the large intestine after the cecum. The intestinal subsegment content was mixed with approximately ∼1 mL of phosphate buffered saline (PBS), removed, and collected. Mucosal washes were collected by gently pipetting ∼1 mL PBS along each subsegment and recovering the solution. Scraping samples were collected by sliding glass microscope slides on the subsegments of the tissue with the mucus, transferring the sample into a tube containing ∼1 mL PBS. All samples were collected in 2 mL tubes, snap frozen in liquid nitrogen and stored at −80°C until further processing.

Human terminal ileum mucosal washes were obtained by colonoscopy using a sterile catheter 230 cm long and 2.3 mm wide (Endo-Flex GmbH). A sterile 10 mL syringe (PosiFlush^TM^ SD) was used to flush physiological saline solution (NaCl 0.9% w/v) onto the mucosal surface in the terminal ileum of the patient. Approximately 1 mL of the resulting intestinal fluid was collected. Biopsies of about 2×2 mm were obtained from the same subsegment using forceps per usual procedural practice. Samples were snap frozen in liquid nitrogen and stored in microcentrifuge tubes at −80°C.

### DNA extraction

Microbial and host (contaminant) DNA from all sample types were extracted using ZymoBIOMICS^TM^ DNA Miniprep Kit (ZYMO Research Europe GmbH, Freiburg, Germany).^17^ The starting material among available for the intestinal contents, mucosal washes, and scraping samples from mice ranged between 24 to 500 mg, 260 to 500 μl, and 13 to 240 mg, respectively. The amounts used for human mucosal washes and whole biopsies ranged from 250 to 600 μl, and 3 to 28 mg, respectively. DNA-/RNA-free water was used as an extraction control.

All samples were mixed with 750 μL of ZymoBIOMICS Lysis Solution, followed by a Proteinase K incubation step using a preheated ThermoBlock at 55°C for 30 min. Bead-beating homogenization was done only for biopsies using a PeQLab Precellys 24 (Bertin Corp., Rockville, MD, USA) for 2 × 15 s at 5500 rpm with beads of 2.0 mm (ZR BashingBead, Lysis Tubes of ZymoBIOMICS). The next steps followed the manufacturer’s protocol. DNA was eluted with 50 μL of DNase/RNase Free Water in a 1.5 mL microcentrifuge tube by centrifuging at 16,000 x g for 3 min. Samples were stored at −80°C until further experiments. DNA quantification and quality check was performed using spectrophotometry in a NanoDrop (PEQLAB Biotechnologie GmbH, ND-1000) and Qubit dsDNA BR Assay Kit or dsDNA HS Assay Kit (invitrogen by Thermo Fisher Scientific).

### Absolute quantification of bacterial load and relative quantification of host DNA

Absolute quantification of the bacterial load was done for all sample types and extraction controls. All amplifications were performed in triplicates in 96-well optical plates (Applied Biosystems) with a final volume of 10 μL containing 5 μL of a 2x SYBR Green PCR Master Mix including a passive reference dye (Applied Biosystems), 10 μM of each primer (Univ337 F 5′-ACT CCT ACG GGA GGC AGC AGT-3′ - Univ 518 R 5′-GTA TTA CCG CGG CTG CTG GCA C-3′ or Univ218 F 5’-ACT GAG ACA CGG CCC A-3’ - Univ515 R: 5’-TTA CCG CGG CMG CTG GCA C-3’) and 1 μL of template DNA (2.5 ng/µL final concentration). Copy numbers per ng of DNA were estimated based on a standard curve prepared with the amplification of the 16S rRNA gene of *E. coli* (27F: 5’-GTT TGA TCC TGG CTC AG-3’ and 1492R: 5’-CGG CTA CCT TGT TAC GAC-3’, by Invitrogen, C404010) in dilutions from 10^3^ to 10^9^. A standard amplification protocol of Applied Biosystems QuantStudio3 AppliedBiosystem (Thermo Fisher Scientific, Darmstadt, Germany) was followed: For samples of mice and humans, amplification was made with 95°C for 20 s followed by 40 cycles at 95°C for 3 s and 60°C per 30 s. In all cases, a melting curve (Tm) analysis was performed, increasing the temperature from 60°C to 95°C at a rate of 0.2°C per second with the continuous monitoring of fluorescence to check for specificity.

Relative quantification of mouse DNA was made using primers (Ms_gDNA_CDC42_F: 5’-CTC TCC TCC CCT CTG TCT TG-3’ and Ms_gDNA_CDC42_R: 5’-TCC TTT TGG GTT GAG TTT CC-3’) for *Mus musculus* nuclear single copy gene *Cdc42*^18^, and SYBR-Green (Applied Biosystems) PCR Master Mix and molecular biology grade water. Amplification conditions for all DNA from mice samples were a denaturation step at 95°C for 2 min, 40 cycles at 95°C for 15 s and 55°C per 15 s. To determine human host DNA content, we amplified a region of the 18S rRNA gene using the primer pairs F: 5’-ACA TCC AAG GAA GGC AGC AG-3’ and R: 5’-TTT TCG TCA CTA CCT CCC CG-3’, amplification conditions were a 95°C for 20 s, 40 cycles at 95°C for 3 s and 60°C per 30 s, with a final melting step.

### Microbial community pre-processing

Amplification of the 16S rRNA gene in the V3-V4 regions was made with Klindworth primers pair (341F: 5’-CCT ACG GGN GGC WGC AG-3’ and 785R: 5’-GAC TAC HVG GGT ATC TAA KCC-3’) from DNA extracted of all the different sample types. Library preparation and sequences were generated by LGC Genomics (LGC Genomics GmbH, Berlin) on the Illumina MiSeq platform in two runs using “v3 chemistry” with 600 cycles (2×300bp). All sequencing raw data can be accessed through the BioProject: PRJNA1043131 in the NCBI Short Read Archive (SRA).

Sequencing reads were processed following filtering by quality check, amplicon sequence variants (ASVs) were inferred and taxonomically assigned using the package DADA2 v1.18.0.^19^ Sequences were trimmed to two different conditions based on the host of origin. For sequences from mice samples, we used the setting truncLen = c(280,210), while for sequences from human samples we used setting truncLen = c(280,230). In both cases we allowed a maximum error of 2 nucleotides and removed the Phix spike-in. Forward and reverse sequencing reads were de-replicated, concatenated and chimeras were removed. Taxonomic assignment was done using the SILVA database (v138.1)^20^ with the RDP naive Bayesian classifier.^21^

The final dataset contained only ASVs assigned at least at the family level, and we removed ASVs classified as mitochondrial or chloroplast. All taxonomic data, ASV abundance table and metadata were compiled into a single object using the package Phyloseq v1.34.0.^22^ A decontamination processing was done using the package Decontam v1.10.0.^23^ We used the combine-either methodology to identify possible bacteria at the genus level classified as contaminants based on their frequency and prevalence with the thresholds (0.1, 0.4, respectively) for mice and (0.2, 0.3, respectively) for human data. Additionally, we included literature research to define the final contaminants, based on whether taxa previously were reported as contaminants or not. ASV tables were filtered out if the samples had less than 100 reads. The read counts were rarefied and transformed to relative abundance for the differential abundance analysis of ASVs at phylum and genus levels.

### Diversity estimation

Beta diversity assessment was based on Aitchison distance of raw counts, using centered log-ratio (CLR) transformation on the data matrix to avoid issues of compositionality.^24^ We applied unconstrained Principal Component Analysis (PCA) to ordinate objects based on such intrasample distances and visualize the multidimensional data, using the vegan R library.^25^ Alpha diversity after rarefaction was evaluated using the Chao1 index for richness taxa estimation, the Shannon index that depends on the richness and evenness of the taxa, and the Inverse Simpson metric, which focuses on a weighted mean of proportional abundances.^26^

### Statistical analysis

Normality was evaluated with the Shapiro test in qualitative variables (DNA concentration, copy number 16S, delta host DNA) using the function shapiro_test() in the R package rstatix (v.0.7.1)^27^, and non-parametric statistics were used when a normal distribution could not be concluded. Comparisons of copy number, host DNA quantity, alpha diversity and differential abundance at different levels of the taxonomy between samples of different types were made using the Kruskal-Wallis test, followed by Dunn’s post hoc test to evaluate per pair of groups in mice samples, and with Mann-Whitney U test in human samples to evaluate the comparisons between two groups. All multiple comparisons were corrected by False Discovery Rate (FDR) with the Benjamini-Hochberg procedure.^28^ The functions used were kruskal_test(), dunn_test() and wilcox_test(), respectively in the rstatix library.^27^

Beta diversity analysis was evaluated throughout PERMANOVA analysis with 999 permutations using adonis2 function in the vegan library (v.2.6-4).^25^ Generalized Linear Mixed Models (GLMMs) were run with the lme4 package function lmer().^29^ To determine the partition of the variance explained in the complex GLMM we used the partR2 package with the function partR2().^30^ The output of this analysis reveals the variance denoted by R^2^ and, the direction and size of the effect indicated by beta. The response variables included quantitative variables as bacterial load, alpha diversity indexes and read counts, and the predictors included the sample types or the genotype or phenotype groups, paired for contrast taking one category as reference. The plots were created with ggplot2().^31^ All the scripts for the analysis described here are available at https://git.bihealth.org/ag-forslund/hydrogels_sampletype.

## Results

We collected intestinal contents from mice in addition to three types of mucus samples, derived from mucosal washes in both hosts, scrapings in mice and biopsies in humans (Supplementary Table S1). For the mice (n = 6) sampling included the collection of three samples each from three different gastrointestinal subsegments per individual, termed scrapings, mucosal washes, and intestinal contents. For the human cohort, we included controls (n = 17) and patients with CD (n = 19), for a total of ten biopsies and 26 washes taken from the terminal ileum (Supplementary Table S1).

### Intestinal mucosal wash sampling captures an equivalent and comparative bacterial load as scraping or biopsy sampling

To determine whether mucosal washes could capture equivalent amounts of bacteria as sampling by scrapings or biopsy, we quantified and observed the resulting DNA concentration and the bacterial load between sample types. In both host species, DNA concentration and bacterial load were significantly different between the sample types (Figure 1a; Supplementary Table S2). Mucosal wash samples exhibited lower DNA concentrations than scrapings or intestinal content samples in mice and biopsies in humans (Figure 1a).

**Figure 1:**
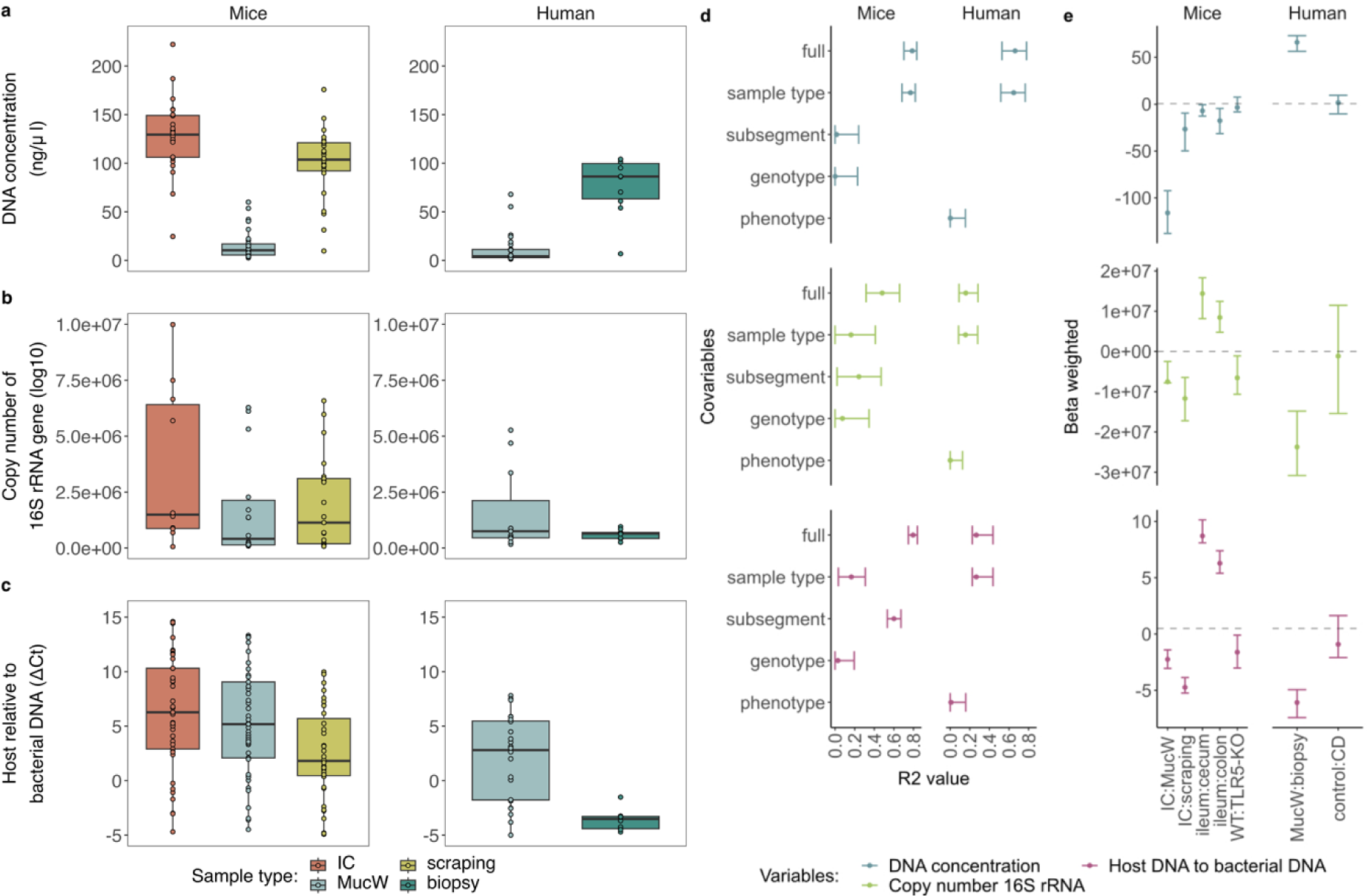
General assessment of bacterial DNA associated with mucus between sample types in mice and humans. In mice and humans, **a)** the DNA concentration (ng/uL), **b)** the bacterial load, assessed as 16S rRNA gene copy number, and **c)** abundance of host DNA relative to bacteria DNA assessed by the ΔCt method. The higher the ΔCt value, the higher the amount of bacteria DNA in a given sample. Each point represents an individual sample. **D)** Variance explained (R^2^) in a forest plot of the DNA concentration, the bacterial load and the host relative to bacterial DNA (ΔCt) in mice (left) and humans (right), with the 95 % confidence interval for each fixed effect in the models with marginal R^2^ values. **e)** Beta estimation of the models per host for DNA concentration (ng/uL), bacterial load and, host DNA relative to bacteria DNA (ΔCt). Margins of the forest plots represent 95% confidence intervals. CD: Crohn’s disease; IC: Intestinal content; MucW: Mucosal washes.

Despite lower overall DNA concentrations, mucosal washes had higher bacterial content than scrapings or biopsy samples (Figure 1b, Supplementary Table S2). Similar results were observed for paired human mucosal washes (n = 9) (Supplementary Figure 1a-b). Additionally, in mice, we compared the bacterial load between the sample types in three subsegments, the ileum, cecum and colon. The bacterial load was similar between mucosal washes and scrapings, but the intestinal content was higher in all subsegments (Supplementary Figure 1b).

To estimate the abundance of host DNA, we used a relative quantification of host DNA genes compared to the bacterial 16S rRNA gene. For mouse samples, we used a *Mus musculus* single-copy nuclear gene *Cdc42,* and for human samples, a region of the 18S rRNA gene. We used ΔCt between host and bacteria as a proxy for host DNA. Thus, the higher the ΔCt value, the higher the content of bacterial DNA in the samples. Mucosal washes yielded less host DNA than scrapings and biopsy samples in mice and humans, respectively (Figure 1c). This was also consistent within subsegments (Supplementary Figure 1c).

To determine the effect of biological covariates (genotype or phenotype, subsegment) on the three variables of interest, we used individual generalized linear mixed models (GLMMs) with DNA concentration, bacterial load and host DNA as our response variables. For mice, the GLMMs included mouse identification number (ID) as random effect, and subsegments, genotypes (WT or TLR5-KO), and sample type (scrapings, mucosal washes, or intestinal contents) as fixed effects. For humans, we included the patient ID as a random effect with phenotype (control or CD) and mucosal wash and biopsy as fixed effects.

We observed that sample type explained most of the variation for the total DNA in both mice (R^2^_marginal_ = 0.770) and humans (R^2^_marginal_ = 0.648), while for the bacterial load it was subsegment in mice (R^2^_marginal_= 0.243) and the sample type in humans (R^2^_marginal_= 0.156). For host DNA, the differences were mostly described by the subsegment in mice (R^2^_marginal_= 0.601) and the sample type in human samples (R^2^_marginal_ = 0.266) (Figure 1d). In all cases, the genotype or phenotype contributed less than 1% of the variation in mice and humans, respectively (Figure 1d; Supplementary Table S3).

The beta estimation of the model was used to determine the direction of the effect. Mucosal wash samples showed a decrease in DNA concentration compared to scraping or biopsy samples in contrast to the intestinal content or the mucosal washes in mice and humans, respectively (Figure 1e). However, mucosal washes and scrapings yielded comparable bacterial loads in mice, and biopsies were lower in human samples. Bacterial DNA relative to host DNA was lower in scrapings and biopsies. On the genotype of TLR5 we observed lower size effect of bacterial load compared to WT mice, and no differentiation on the phenotype between all controls and CD patients on DNA concentration, bacterial load, or host DNA (Figure 1e).

Furthermore, pairwise contrasts of the bacterial load between the sample types within the three subsegments were statistically different only for intestinal content and scraping sample types, with an increase for intestinal content (Supplementary Table S4). For human samples, no differences in bacterial load were observed when contrasting sample type with phenotype groups (Supplementary Table S4).

The results suggest that intestinal washes are contaminated with less host DNA than scrapings and biopsies, a desirable feature to avoid wasting bacterial resolution by sequencing host DNA in the samples. Bacterial load is comparable between the mucosal washes and scrapings in mice, but still it is lower in biopsies compared to mucosal washes in humans.

### Mucosal wash samples yielded less overall sequenced DNA but also less mitochondrial contamination than scraping and biopsies

To understand the impact of sample type on sequencing results, we quantified its effect on total reads as well as reads remaining after taxonomy filtering and decontamination using the Decontam v1.10.0 package.^23^ We identified a total of 3 and 16 specific contaminants in mice and humans, respectively (Supplementary Figures 2a-f). Of these, one mouse contaminant (*Cutibacterium*) and two human contaminants (*Delftia, Sphingomonas*) were found in the extraction controls. We excluded ASVs previously classified as contaminants by the Decontam library but known to inhabit the host intestine, otherwise they were discarded as contaminants (Supplementary Table S5). ASVs classified as *Cutibacterium* were also excluded from human samples as they had a prevalence of 17%.

The number of total mitochondrial-associated and decontaminated bacterial reads was lower in mucosal wash samples compared to scrapings in mice, with similar results per subsegment (Supplementary Figure 3a-c). Conversely, in humans, mucosal washes had higher values for all the read variables (Figure 2a-c; Supplementary Figure 3a-c). The evaluation with the remaining covariates showed that the variance for the total and filtered reads in mice was explained by the subsegments (R^2^_marginal_= 0.182 and R^2^_marginal_= 0.222), and the mitochondrial reads were mainly explained by the genotype (R^2^_marginal_= 0.053) in mice, while in humans all read variables were mostly explained by the sample type (R^2^_marginal_= 0.206, R^2^_marginal_= 0.242, R^2^_marginal_= 0.048, respectively) (Figure 2d and Supplementary Table S3). We also observed that mucosal wash samples generally yielded fewer total or filtered reads than intestinal content samples, and the genotype TLR5 had a higher effect on the filtered reads compared to WT mice. Interestingly, total or filtered reads had a lower effect from biopsies in contrast to mucosal washes in human samples, and a similar effect between both phenotypes in the three read variables were obtained (Figure 2e and Supplementary Table S3). However, no pairwise contrast between sample types within genotypes or subsegments reached significance in mice, and humans samples had a significant difference on total and filtered reads when comparing the mucosal washes and biopsies stratifying only within CD patients or control patients (Supplementary Table S4).

**Figure 2:**
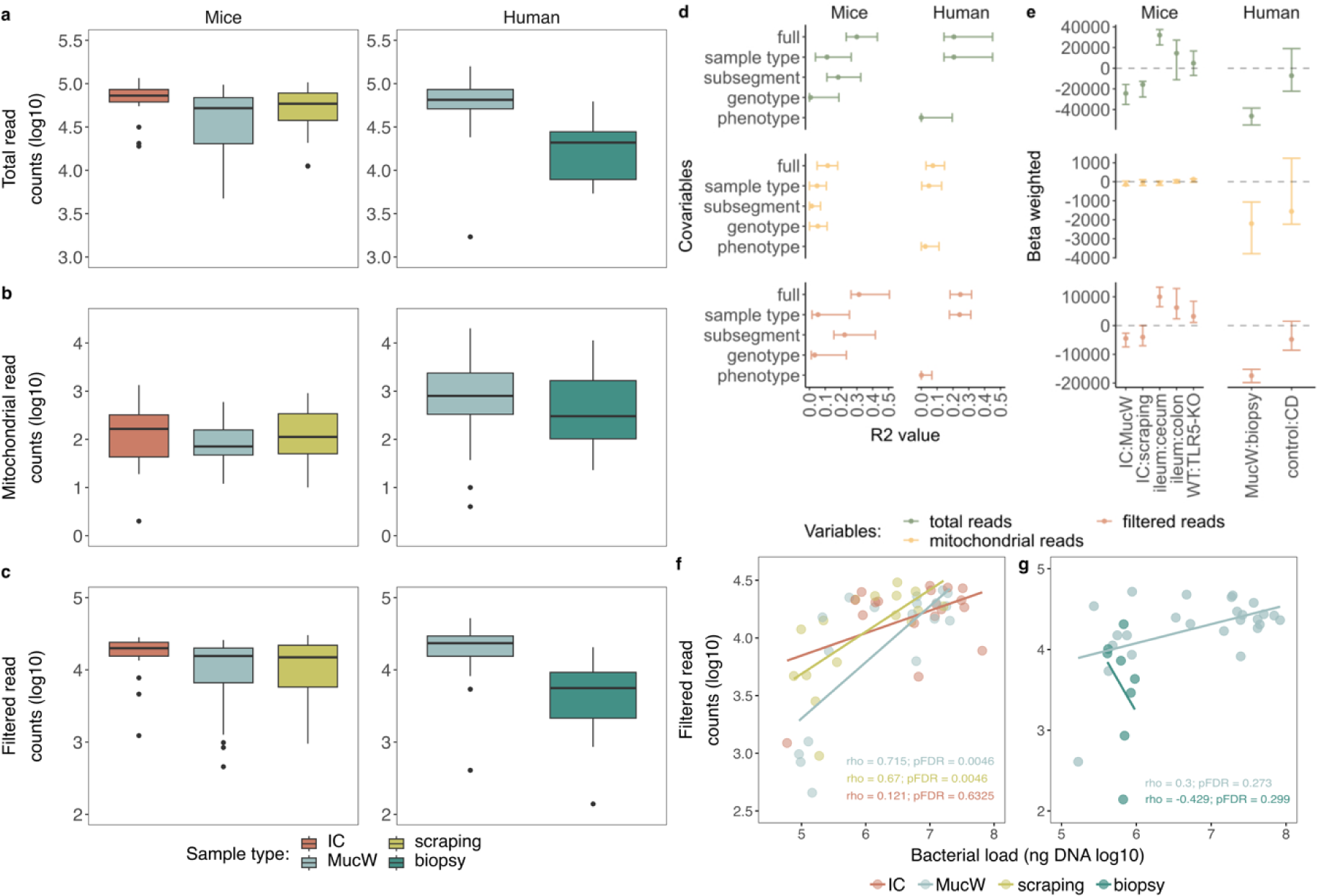
Assessment of sequencing reads between sample types in mice and humans. **a)** Total read counts, **b)** mitochondrial reads, **c)** filtered read counts, represented in logarithm 10**. d)** Variance explained (R^2^) in a forest plot of the total, mitochondrial and filtered reads in mice (left) and humans (right), with the 95% confidence interval for each fixed effect in the models with marginal R^2^ values. **e)** Beta estimation of the models per host mice (left) and human (right). Margins of the forest plots represent 95% confidence intervals. **f-g)** Spearman correlation between the bacterial load (ng of DNA represented in logarithm 10) and the filtered read counts (represented in logarithm 10) in **f)** mice and **g)** humans. IC: Intestinal content; MucW: Mucosal washes.

We evaluated whether the bacterial load was associated with the decontaminated bacterial reads in the samples. We performed a Spearman correlation between the two proxies. Higher bacterial load was strongly correlated with higher decontaminated bacterial reads in mucosal wash and scraping samples from mice (⍴ > 0.6, pFDR < 0.01). However, we did not observe a significant correlation of these parameters in human samples (Figure 2f-g).

### Diversity and composition are comparable between intestine mucosal washes, scraping and biopsy samples

To assess intra-individual microbial diversity, we calculated the Chao1 index for ASV richness estimation, Shannon, and Inverse Simpson indices. Notably, we did not observe statistically significant differences between sample types in either mouse or human samples, suggesting comparable diversity readouts from the different sample types. Nevertheless, it is relevant to acknowledge that larger sample sizes would allow us to detect more subtle variation.

We applied GLMM models to investigate the factors influencing alpha diversity metrics and to observe the effect size in mice and humans. Alpha diversity metrics are essential for assessing species diversity within a community. For mice, our analysis revealed that the primary factors influencing the variation in Chao1 and Shannon diversity are related to the subsegment (Chao1: R^2^_marginal_ = 0.501; Shannon: R^2^_marginal_ = 0.306). Additionally, the mouse genotype was found to have a significant effect impact on species evenness (R^2^_marginal_ = 0.213), suggesting the influence of the disease genotype on the less dominant bacterial species. A higher effect on Chao1 was observed from the mucosal washes but lower on Shannon for the scraping and mucosal washes contrasting to the intestinal content, and a similar effect was observed for between the genotype groups in all alpha diversity indexes.

In the human dataset, the primary driver of variation in alpha diversity metrics was the phenotype, specifically for Shannon and species evenness (Shannon: R^2^_marginal_ = 0.021; Inverse Simpson: R^2^_marginal_ = 0.017). A lower effect size on Chao1 and Shannon was observed on the biopsies compared to the mucosal washes and, on the patients with CD to the controls (Figure 3e). Our pairwise comparisons revealed no statistically significant differences in alpha diversity metrics between groups for either mouse or human samples (Supplementary Table S4). These results provide insight into the key factors that may influence alpha diversity in mice and humans, as impact of subsegment and genotype predominate in mice and sample type and phenotype in humans.

**Figure 3:**
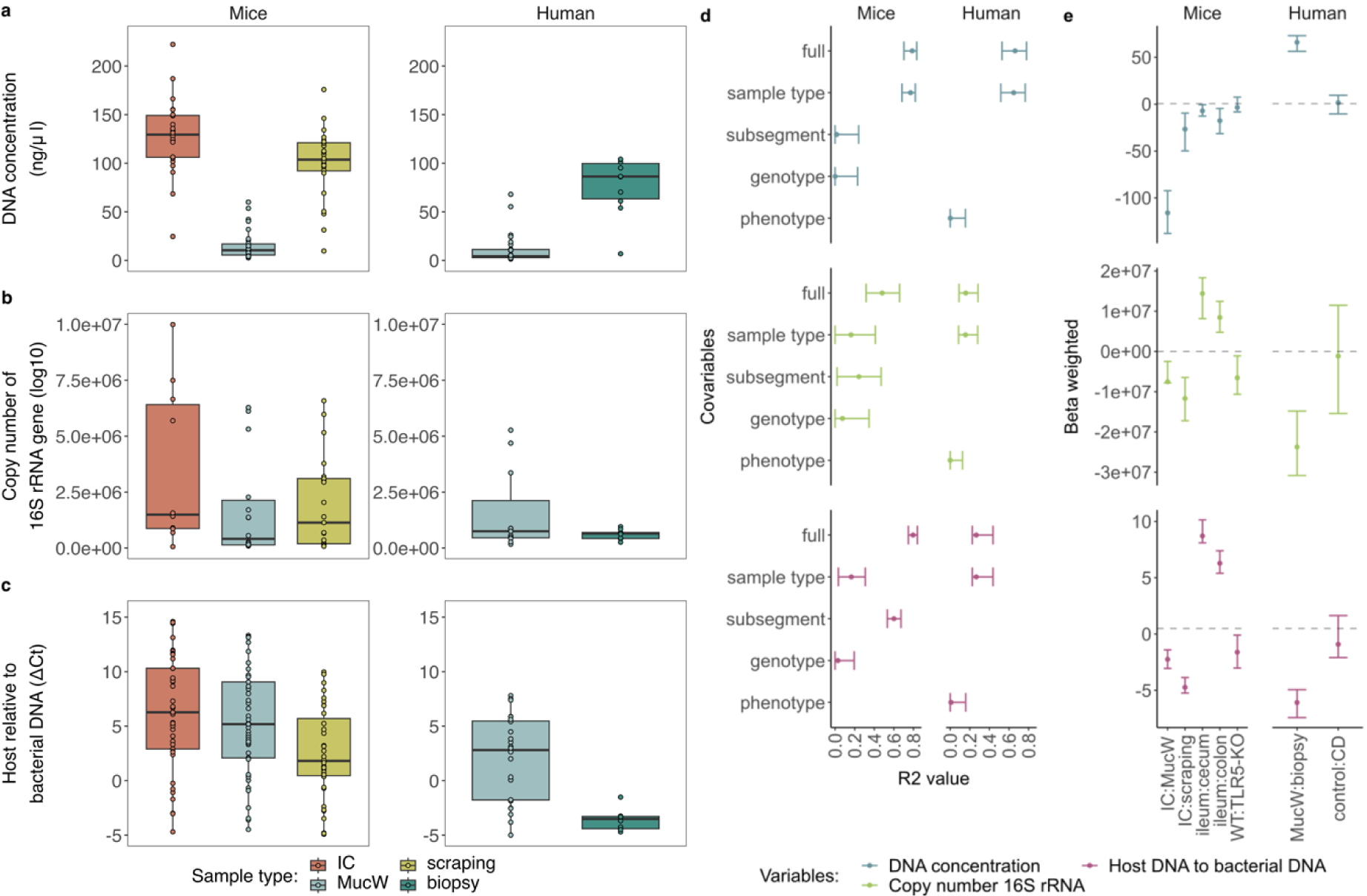
Assessment of alpha and beta diversity between sample types in mice and humans. Alpha diversity is presented with the metrics: **a)** Chao1, **b)** Shannon, **c)** inverse Simpson in mice (left) and humans (right). **d)** Variance explained (R^2^) in a forest plot of the alpha diversity metrics, with the 95% confidence interval for each fixed effect in the models with marginal R^2^ values in mice (left) and humans (right). **e)** Beta estimation of the models per host mice (left) and human (right). Margins of the forest plots represent 95% confidence intervals. **f-g)** Principal Component Analysis (PCA) of the beta diversity represented with Aitchison distance in **f)** mice and **g)** humans. IC: Intestinal content; MucW: Mucosal washes.

Changes in bacterial composition due to beta diversity were determined using the Aitchison distance. Only for the human samples we observed that the structure of the bacterial community was variable on the x-axis with a 13.696 %, while the mouse samples were more homogeneously distributed (Figure 3f-g). Permutational analysis showed that the genotype groups significantly explained over 13 % of the variance in mouse bacterial composition. Additionally, the sample type was not a significant predictor of overall compositional variability in mice. In human samples, the bacterial composition was significantly explained by the sample type, and the phenotype with a lower variance, corresponding to 28.96 % and 4.97 %, respectively (Table 1).

**Table 1:**
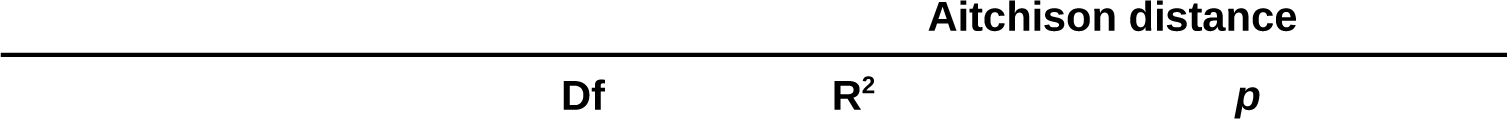
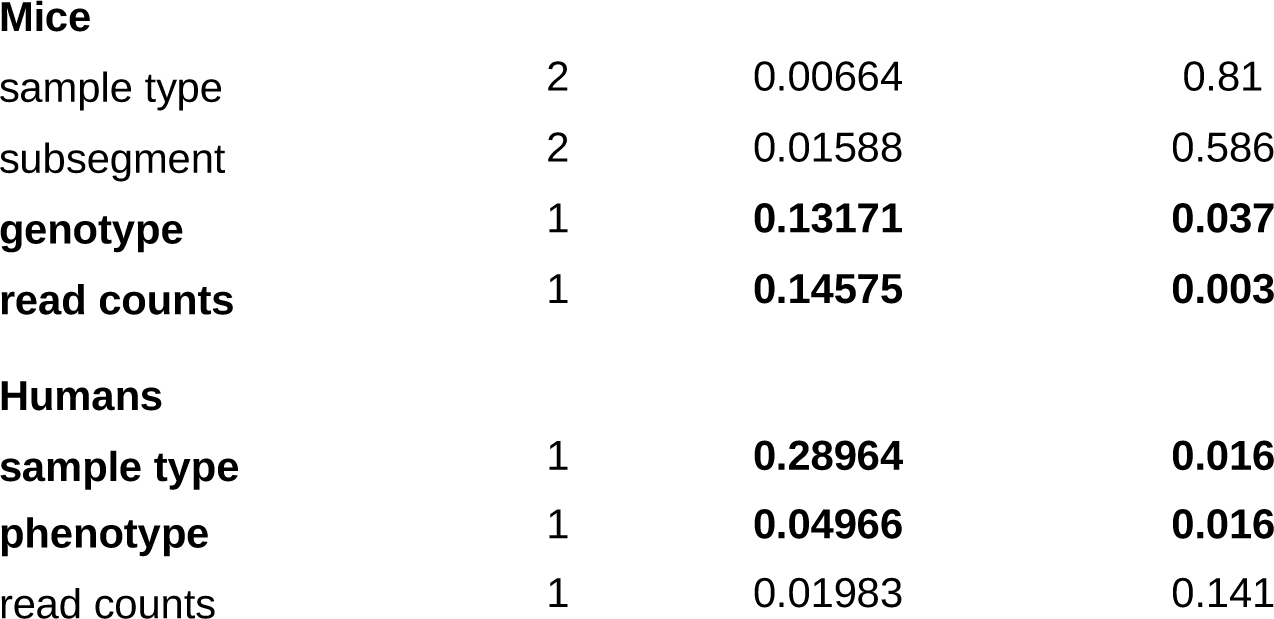
Permutational analysis of variance of the beta dispersion for bacterial taxa composition in mice and human GI samples.

| Aitchison distance |  |  |
| --- | --- | --- |
| Df | $R^2$ | $p$ |

Mice
|  |  |  |  |
| --- | --- | --- | --- |
| sample type | 2 | 0.00664 | 0.81 |
| subsegment | 2 | 0.01588 | 0.586 |
| **genotype** | **1** | **0.13171** | **0.037** |
| **read counts** | **1** | **0.14575** | **0.003** |

Humans
|  |  |  |  |
| --- | --- | --- | --- |
| sample type | 1 | 0.28964 | 0.016 |
| **phenotype** | **1** | **0.04966** | **0.016** |
| read counts | 1 | 0.01983 | 0.141 |

To determine whether taxonomic differences were observed between sample types, we compared the relative abundance of bacteria at the phyla and genus taxonomic levels. In mice, we observed high similarity between sample types when comparing at the phyla level, especially for the dominant Firmicutes and Bacteroidota (Supplementary Figure 4a-b). When comparing abundances clustered at the genus level, we observed that all genus abundances were comparable between the sample types (Figure 4a). In human samples, we also observed no significant change in abundance between sample types at the phyla or genus level (Figure 4b, Supplementary Figure 5a-b).

**Figure 4:**
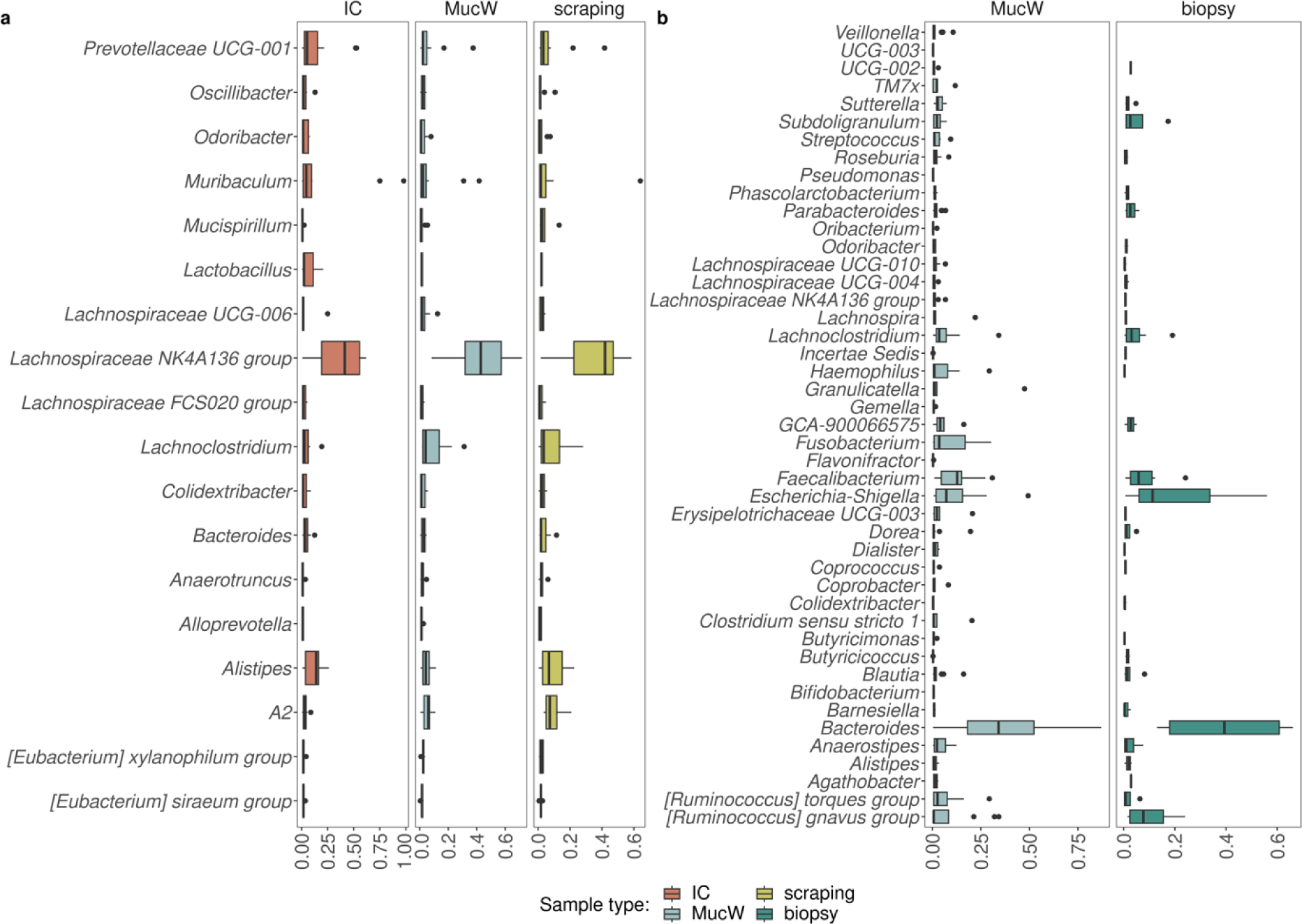
Evaluation of the relative abundance at genus level between sample types per host. Abundance was assessed between sample types in **a)** mice (left) and **b)** humans (right). The x-axis represents the relative abundance, and on the y-axis the genus present in the samples with a frequency greater than 4 per host. IC: Intestinal content; MucW: Mucosal washes.

## Discussion

Numerous studies have highlighted the relevance of assessing mucus-associated microbiota in host organisms.^5^ Our investigation focused on mice and humans (the primary preclinical and clinical settings, respectively), using samples previously used for mucus-associated microbiota studies. Notably, our results demonstrate that mucosal wash samples are a viable alternative to biopsy in humans or scraping in mice, providing a simpler and less invasive option while maintaining the transferability of results.

We employed quantitative PCR to determine 16S rRNA gene copies per sample.^32^ As expected, intestinal contents yielded higher bacterial loads than scrapings and mucosal wash samples, particularly for mouse cecum and colon. The latter shows a higher bacterial cell count in the mouse colon than in the small intestine, consistent with previous observations.^15^ Notably, the mucosal wash samples yielded similar amounts of bacterial DNA as scraping samples, despite the difference in total DNA extracted. Our results further confirm that the mucosal wash method of sample collection is effective for obtaining bacterial DNA from the mucus.

Mucosal wash samples have previously been used for microbiome assessment, with suggestions that the resulting higher bacterial DNA content may be of benefit for low-biomass samples.^10,12,15^ Our study shows that mucosal washes not only yield higher bacterial loads, but also significantly lower host DNA content compared to scraping samples in mice or biopsies in humans, as assessed by relative quantification of 16S rRNA and 18S rRNA, particularly in the cecal subsegment. Our findings confirm that most of the DNA in biopsies and scrapings is of host origin.

We sought to confirm whether the higher proportion of host DNA affected sequencing performance by comparing total reads, mitochondrial assigned reads, and bacterial reads. While our computational analysis indicated lower raw mitochondrial reads in mouse mucosal wash samples, this trend was not replicated in human samples, with no variance in mitochondrial reads between sample types. However, total and filtered bacterial reads were higher in mucosal washes than in human biopsies. This is supported by previous reports that have shown that human biopsies had more mitochondrial and chimeric reads, compared to lavages or brushing samples.^12^

Contrary to previous studies demonstrating higher richness estimates (Chao1) and Shannon diversity in the lumen than in the mucosa of mice^15,33^, our results showed similar values in the luminal intestinal content to either mucosal washes or scraped mucus samples in mice. In humans, comparisons of microbiota alpha diversity between sample types in the terminal ileum showed that mucosal washes had greater richness than biopsies. Our findings support previous reports suggesting lower microbial biomass in biopsies than in aspirates or lavages.^10,11^

We found that for beta diversity in mouse samples, the sample type did not significantly influence the overall composition of the bacterial communities. Instead, our analysis revealed that most of the composition variance was explained by the different genotypes among the mice. In contrast, in humans the sample type and the phenotype (disease) groups emerged as significant predictors of beta diversity.

When collecting mouse mucosal samples, we observed that using a mucosal wash method instead of scraping had a minimal effect on the overall composition of the bacterial communities. This suggests that the choice between mucosal washes and scraping does not significantly impact the estimated microbial composition of mouse samples.

Firmicutes and Bacteroidota predominated among mucus-associated bacteria in both host models, followed by Proteobacteria in human ileal biopsies and mucosal wash samples. Our observation is consistent with similar studies evaluating colon biopsies by 16S rRNA or shotgun sequencing.^14^ Surprisingly, we did not observe the enrichment of *Ruminococcaceae* the three different subsegments in mouse samples or *Lachnospiraceae* in human ileum samples, despite their typical association with colonic mucus-associated microbiota.^34^ However, one genus *Lachnospiraceae*, part of the Firmicutes phylum, was present in all mouse sample types. Overall, mucosal washes are compositionally similar to the other types tested but have the added benefit of relatively low host contamination.

Our study is the first to compare the mucus-associated microbiota in biopsies obtained from mucosal wash samples of the human terminal ileum. Our findings reveal that the composition of mucus-associated microbiota from the terminal ileum is not strongly affected by the choice between these two sampling protocols, but with mucosal wash samples recovering a greater proportion of bacterial reads. This supports the feasibility of using mucosal washes from the ileum or colon as a reproducible surrogate for colonic mucosal biopsies.^11^ Importantly, we observed similar microbiota compositions in mucosal washes and biopsy samples, suggesting their interchangeable use for future experiments assessing mucosa-associated microbiota.

### Conclusions

In conclusion, our comprehensive assessment of mucus-associated microbiota in both mice and humans has provided compelling evidence for the suitability of mucosal wash samples as a robust alternative to traditional biopsies or other tissue-derived samples such as scrapings. This study not only confirms the comparable nature of the microbial composition recovered by these sampling protocols but also highlights the distinct advantage of mucosal wash samples - their higher bacterial content and lower host DNA content. These findings underscore the potential for reproducible and minimally invasive techniques in the field of mucus-associated microbiota research.

## Acknowledgments

This study was funded by the Deutsche Forschungsgemeinschaft (DFG) Collaborative Research Center (SFB) 1449: Dynamic Hydrogels at Biointerfaces, Project ID 431232613 (to MF, SKF, BS); TRR241 project-ID 37587604 [BS] and CRC 1449 project ID 431232613 (BS).

## Data availability statement

**Data deposition**

**BioProject ID:** PRJNA1043131

