## Supplementary Figure for "Mucosal washes are useful for sampling intestinal mucus-associated microbiota despite low biomass"

**Supplementary Figure 1:** **a)** DNA concentration, **b)** bacterial load (copy number of 16S rRNA gene in log10), and **c)** host relative to bacterial DNA (ΔCt) between samples within subsegments in mice (n = 6 per subsegment) and in the paired washing-biopsy human samples (n = 9).

**Supplementary Figure 2: a-b)** Scatterplots with histogram representing the p-scores generated with the combined-either method based on frequency (x-axis) and prevalence (y-axis) using the Decontam library. Colours represent the ASVs classified as true (yellow) or false (blue) contaminants, and bars represent the distribution of the ASVs. **a)** mice samples, **b)** human samples. **c** and **e)** Lollipop graph representing the frequency based on the abundance of the ASVs classified as contaminants, y-axis names are a combination of the family and genus, and the colour of the point represents the phylum to which the ASV belongs, **c)** mice samples, **e)** human samples. **d** and **f)** Lollipop graph representing the prevalence of the ASVs classified as contaminants, y-axis names are a combination of the family and genus, and the colour of the point represents the phylum to which the ASV belongs. **d)** mouse samples, **f)** human samples.

**Suppl****ementary Figure 3:** **a)** Total read counts, **b)** mitochondrial read counts, and **c)** filtered read counts represented in logarithm 10 between samples within subsegments in mice (n = 6 per subsegment) and in the paired washing-biopsy human samples (n = 7).

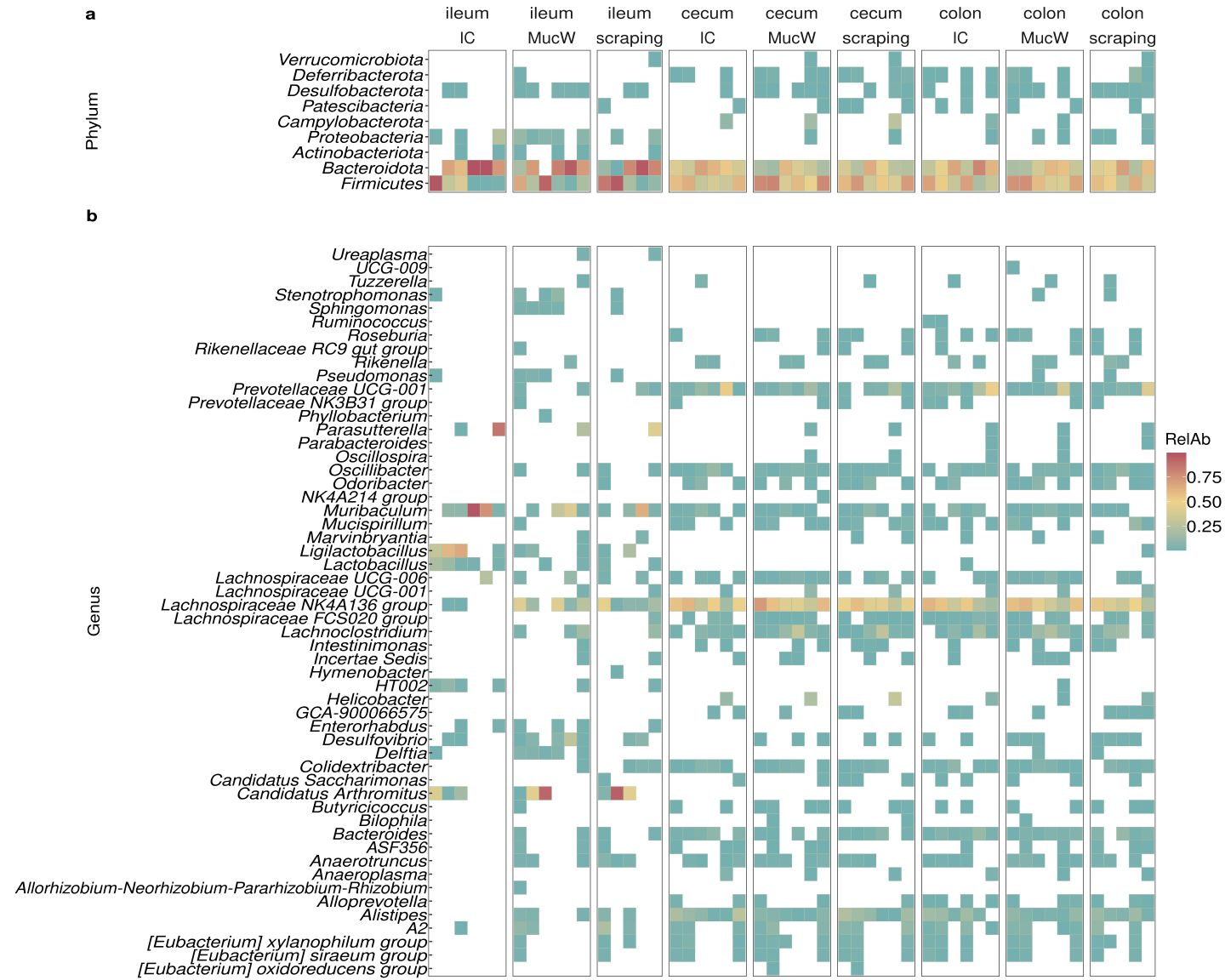

**Supplementary Figure 4:** **a-b)** Heatmap of the relative abundance of the phyla (y-axis) present per sample (x-axis) and grouped by subsegment and mice sample types. **c-d)** Heatmap of the relative abundance of the phyla (y-axis) presented per sample (x-axis) and grouped by human sample types.

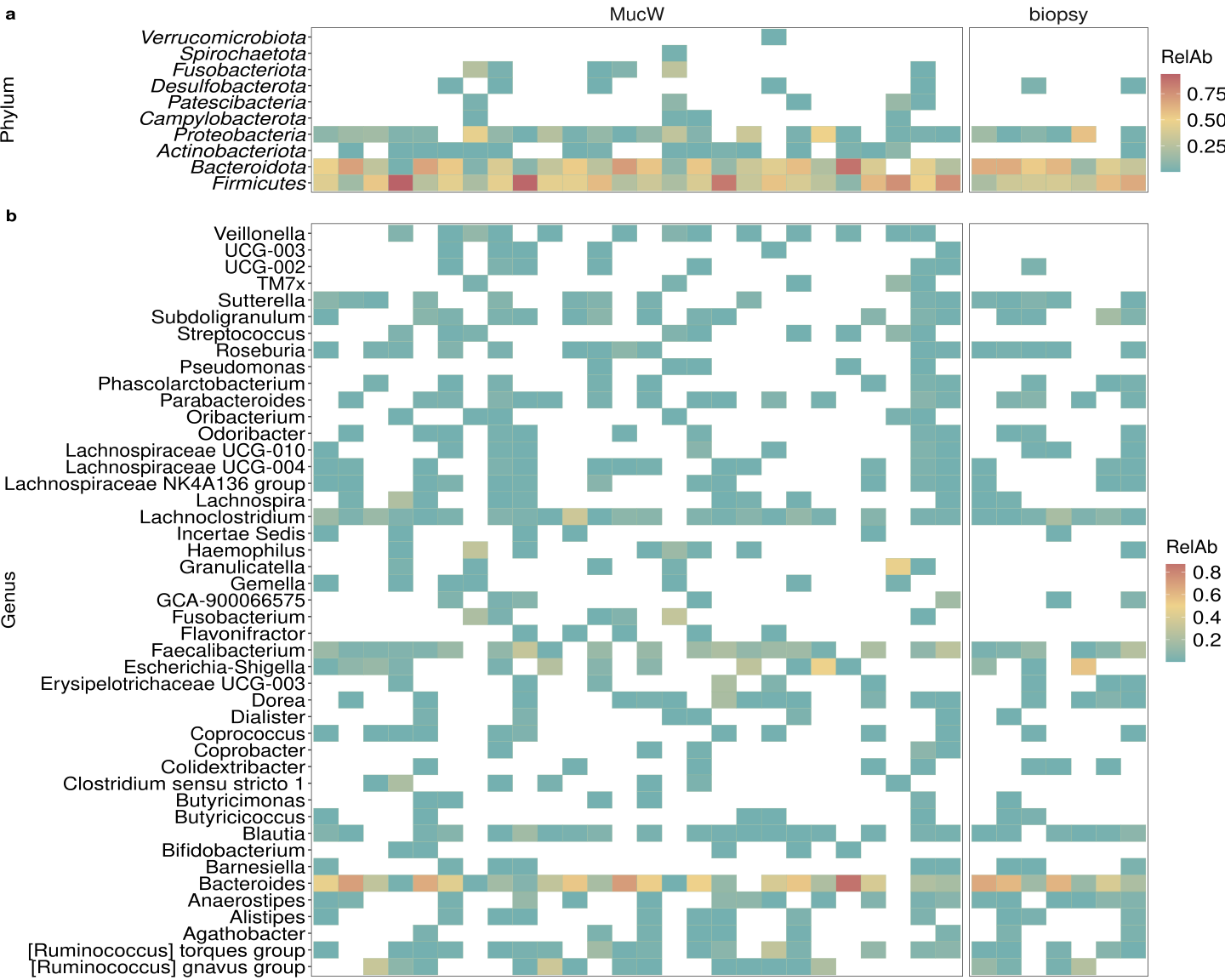

**Supplementary Figure 5:** **a-b)** Heatmap of the relative abundance of the phyla (y-axis) presented per sample (x-axis) and grouped by human sample types.
